# Graph neural fields: a framework for spatiotemporal dynamical models on the human connectome

**DOI:** 10.1101/2020.09.08.287110

**Authors:** Marco Aqil, Selen Atasoy, Morten L. Kringelbach, Rikkert Hindriks

**Author notes:** Spinoza Centre for Neuroimaging, Amsterdam, The Netherlands (Marco Aqil).

## Abstract

Tools from the field of graph signal processing, in particular the graph Laplacian operator, have recently been successfully applied to the investigation of structure-function relationships in the human brain. The eigenvectors of the human connectome graph Laplacian, dubbed “connectome harmonics”, have been shown to relate to the functionally relevant resting-state networks. Whole-brain modelling of brain activity combines structural connectivity with local dynamical models to provide insight into the large-scale functional organization of the human brain. In this study, we employ the graph Laplacian and its properties to define and implement a large class of neural activity models directly on the human connectome. These models, consisting of systems of stochastic integrodifferential equations on graphs, are dubbed *graph neural fields*, in analogy with the well-established continuous neural fields. We obtain analytic predictions for harmonic and temporal power spectra, as well as functional connectivity and coherence matrices, of graph neural fields, with a technique dubbed CHAOSS (shorthand for *Connectome-Harmonic Analysis Of Spatiotemporal Spectra*). Combining graph neural fields with appropriate observation models allows for estimating model parameters from experimental data as obtained from electroencephalography (EEG), magnetoencephalography (MEG), or functional magnetic resonance imaging (fMRI); as an example application, we study a stochastic Wilson-Cowan graph neural field model on a high-resolution connectome, and show that the model equilibrium fluctuations can reproduce the empirically observed harmonic power spectrum of BOLD fMRI data. Graph neural fields natively allow the inclusion of important features of cortical anatomy and fast computations of observable quantities for comparison with multimodal empirical data. They thus appear particularly suitable for modelling whole-brain activity at mesoscopic scales, and opening new potential avenues for connectome-graph-based investigations of structure-function relationships.

**Author summary:** The human brain can be seen as an interconnected network of many thousands neuronal “populations”; in turn, each population contains thousands of neurons, and each is connected both to its neighbors on the cortex, and crucially also to distant populations thanks to long-range white matter fibers. This extremely complex network, unique to each of us, is known as the “human connectome graph”. In this work, we develop a novel approach to investigate how the neural activity that is necessary for our life and experience of the world arises from an individual human connectome graph. For the first time, we implement a mathematical model of neuronal activity directly on a high-resolution connectome graph, and show that it can reproduce the spatial patterns of activity observed in the real brain with magnetic resonance imaging. This new kind of model, made of equations implemented directly on connectome graphs, could help us better understand how brain function is shaped by computational principles and anatomy, but also how it is affected by pathology and lesions.

## Introduction

The spatiotemporal dynamics of human resting-state brain activity is organized in functionally relevant ways, with perhaps the best-known example being the “resting-state networks” [1]. How the repertoire of resting-state brain activity arises from the underlying anatomical structure, i.e. the connectome, is a highly non-trivial question: it has been shown that structural connections imply functional ones, but that the converse is not necessarily true [2]; furthermore, specific discordant attributes of structural and functional connectivity have been found by network analyses [3]. Research on structure-function questions can be broadly divided into data-driven (analysis), theory-driven (modelling), and combinations thereof. In this work, we combine techniques from graph signal processing (analysis) and neural field equations (modelling) to outline a promising new approach for the investigation of whole-brain structure-function relationships.

A recent trend of particular interest in neuroimaging data analysis is the application of methods from the field of graph signal processing [4–8]. In these applications, anatomical information obtained from diffusion tensor imaging (DTI) and structural MRI is used to construct the *connectome graph* [9], and combined with functional imaging data such as BOLD-fMRI or EEG/MEG to investigate structure-function relationships in the human brain (see [10, 11] for reviews). The workhorse of graph signal processing analysis is the *graph Laplacian operator*, or simply graph Laplacian. Originally formulated as the graph-equivalent of the Laplace-Beltrami operator for Riemannian manifolds [12, 13], the graph Laplacian is now established as a valuable tool in its own right [10]. The eigenvectors of the graph Laplacian provide a generalization of the Fourier transform to graphs, and therefore also a complete orthogonal basis for functions on the graph. In the context of the human connectome graph, the eigenvectors of the graph Laplacian are referred to as *connectome harmonics* by analogy with the harmonic eigenfunctions of the Laplace-Beltrami operator. Of relevance to the current work, several connectome harmonics have been shown to be related to specific resting-state networks [9]. More recent studies have provided additional evidence for this claim [14], and others used a similar approach to explain how distinct electrophysiological resting-state networks emerge from the structural connectome graph [15]. Furthermore in [9], for the first time, to the best of our knowledge, a model of neural activity making use of the graph Laplacian was implemented, and used to suggest the role of excitatory-inhibitory dynamics as possible underlying mechanism for the self-organization of resting-state activity patterns. In other very recent work [16] graph-Laplacian-based techniques were employed to model MEG oscillations. Considering these developments, the combination of neural activity modelling and graph signal processing techniques appears as a promising direction for further inquiry.

Whole-brain models are models of neural activity that are defined on the entire cortex and possibly on subcortical structures. This is generally achieved either by parcellating the cortex into a network of a few dozens of macroscopic, coupled regions of interest (ROIs), or by approximating the cortex as a bidimensional manifold, and studying continuous integrodifferential equations in a flat or spherical geometry (See [17] for a review). In this study, relying on graph signal processing methods such as the graph Laplacian and graph filtering [6, 8], we show how to define and implement a large class of whole-brain models of neural activity on arbitrary metric graphs^1^, and in particular on a non-parcellated mesoscopic human connectome. These models consist of systems of integrodifferential equations, and are dubbed *graph neural fields* by analogy with their continuous counterparts. We obtain analytic expressions for harmonic and temporal power spectra, as well as functional connectivity and coherence matrices, of graph neural fields, with a technique dubbed CHAOSS (shorthand for *Connectome-Harmonic Analysis Of Spatiotemporal Spectra*). When combined with appropriate observation models, graph neural fields can be fitted to and compared with functional data obtained from different imaging modalities such as EEG/MEG, fMRI, and positron emission tomography (PET). As an example application, we study a Wilson-Cowan stochastic graph neural field model, implemented on a single-subject unparcellated connectome. We show that the model can accurately reproduce the empirical harmonic spectrum of resting-state BOLD signal fluctuations. In sum, graph neural fields provide a computationally efficient and versatile modelling framework that is tailored for connectome-graph-based structure-function investigations, and particularly suitable for modelling whole-brain activity on mesoscopic scales. Graph neural fields present immediate application in the investigation of the relationship between individual anatomy, pathology, and lesions with functional activity; and furthermore provide a model-based approach to test novel graph signal processing neuroimaging hypotheses and analyses.

In Section *Results* we start by providing an analytic solution and a numerical implementation of the damped-wave equation on the human connectome graph, since this equation is of interest in the context of modelling neural activity propagation. Next, we show how to implement the Wilson-Cowan stochastic graph neural field model on arbitrary metric graphs. We obtain results of linear stability analysis, CHAOSS, and numerical simulations, first on a one-dimensional graph with 1000 vertices, and then on a single-subject connectome consisting of approximately 18000 cortical vertices and 10000 white matter tracts. The simplified context of a 1-dimensional graph is useful to study the effect of simple graph properties, such as inter-node distance and long-range connections, on model dynamics; moving to a real-world application, we fit the full-connectome model parameters to the experimentally observed harmonic power spectrum of resting-state fMRI data of a single subject, showing excellent agreement between the analytically predicted, numerically simulated, and empirical power spectra. In Section *Methods* we provide a more general and detailed description of the framework of graph neural fields. We define spatiotemporal convolution on graphs through the weighted graph Laplacian, and obtain the graph-equivalents of several connectivity kernels and reaction-diffusion systems that are of interest for neural activity modelling. Finally, we show how to derive analytic expressions for harmonic and temporal power spectra, as well as coherence and functional connectivity matrices, of graph neural fields (CHAOSS).

## Results

### Damped wave equation on the human connectome graph

The damped-wave equation describes the dynamics of simultaneous diffusion and wave propagation, and is thus of interest in the context of modelling activity propagation in neural tissue [18]. Here, we solve the graph equivalent of the damped-wave equation and implement it on the human connectome graph. In one-dimensional continuous space, the damped-wave equation is

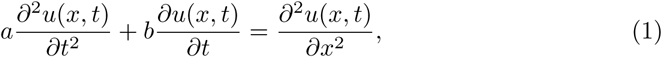

where *a* and *b* are scalar parameters. Its graph-equivalent is given by

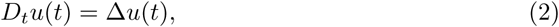

where *u*(*t*) is a function on the graph, Δ is the distance-weighted graph Laplacian, and *D*_*t*_ = *ad*^2^*/dt*^2^ + *bd/dt*. Since the graph Laplacian is a constant matrix, we can straightforwardly obtain the exact solution at time *t*:

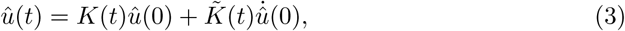

where 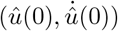 are the initial conditions, the superscript *û* indicates the graph Fourier transform (see *Methods*), and

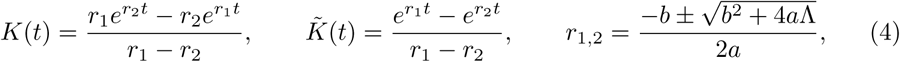

Λ is the diagonal matrix of graph Laplacian eigenvalues. Having obtained an exact solution, we can efficiently simulate the time-evolution of the damped-wave equation on arbitrary metric graphs, for example with the following numerical scheme:

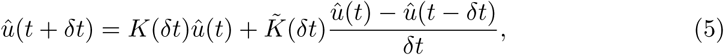

We also note that the Telegrapher’s Equation, which is of interest in the context of modelling action potentials [19]

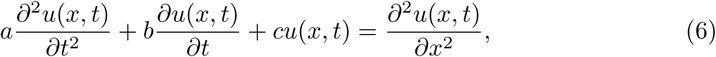

can also be implemented on metric graphs simply by substituting Λ with (Λ − Diag(*c*)) in Eq (4).

Fig 1 shows an example implementation of Eq (5) on the human connectome graph. The initial condition is a Gaussian centered in the occipital cortex of the left hemisphere, itself obtained by applying the graph Gaussian convolution filter (Table 2) to an impulse function. Note how the initially localized activity propagates with a characteristic speed and wavelength. Damping of the wave, caused by the diffusion term, can also be observed. Fig 2 shows an implementation with different parameters, giving rise to faster wavefront propagation and less damping. Fig 3 shows the same implementation as Fig 2, from a different point of view and with a narrower color-scale. This is to emphasize that, aside from the dominant surface-based wavefront, activity also propagates non-locally along white-matter fibers.

**Table 1.**
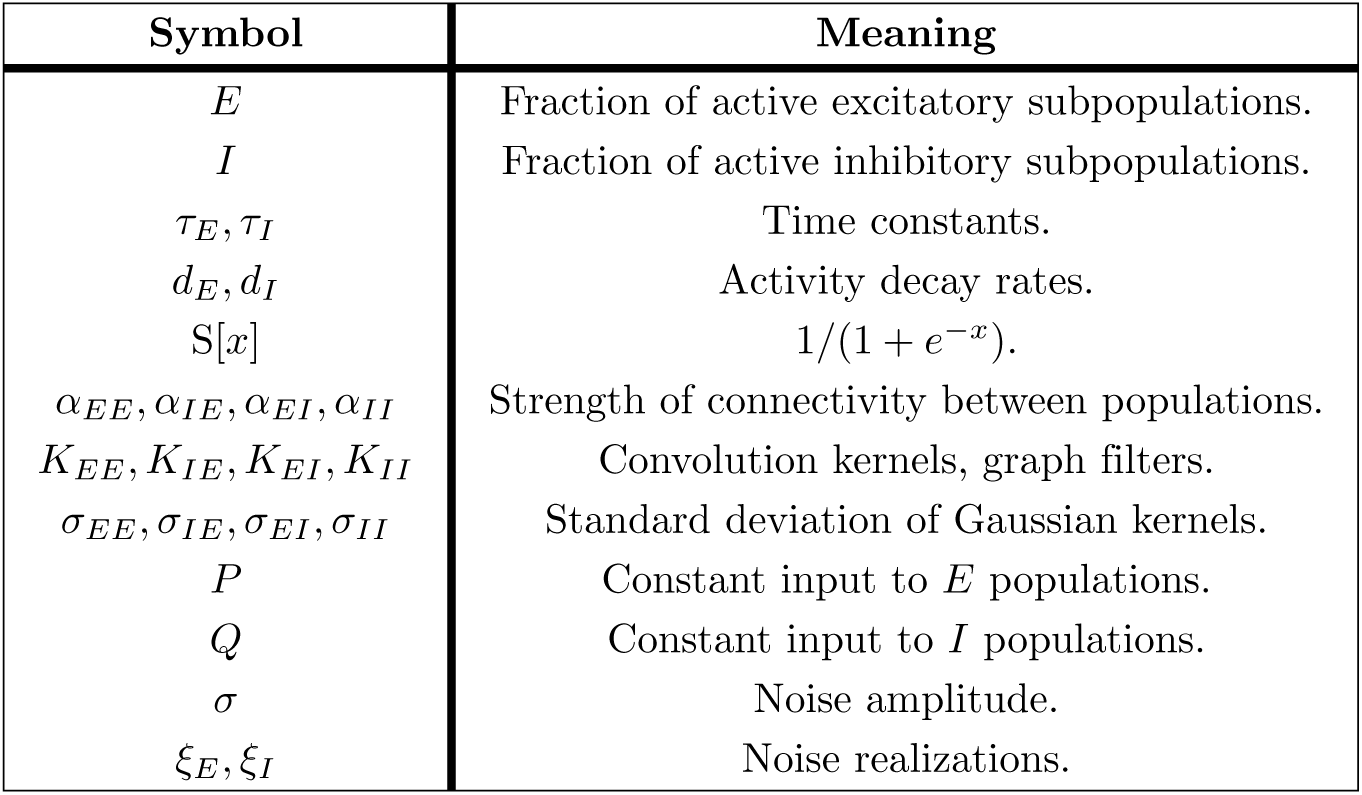
Meaning of symbols in the continuous Wilson-Cowan equations and their graph equivalents.

**Table 2.**
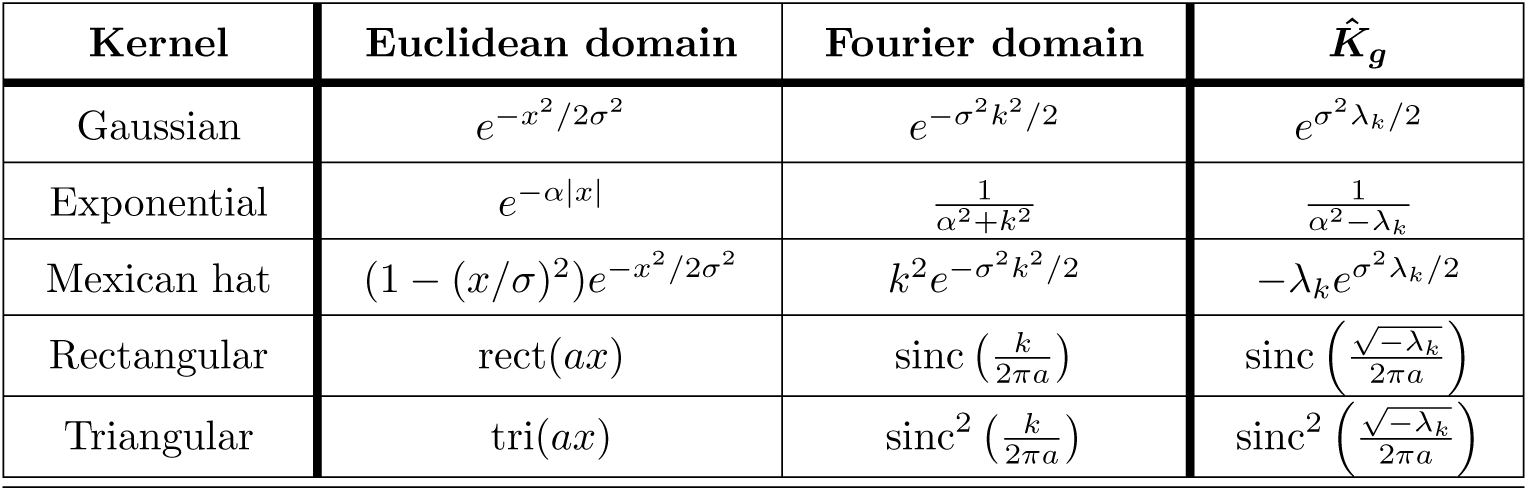
Spatiotemporal convolution kernels in the Euclidean, Fourier, and graph domains. Normalization factors are omitted. The sampling function sinc is defined as sinc(*x*) = sin (*πx*)*/*(*πx*).

**Fig 1.**
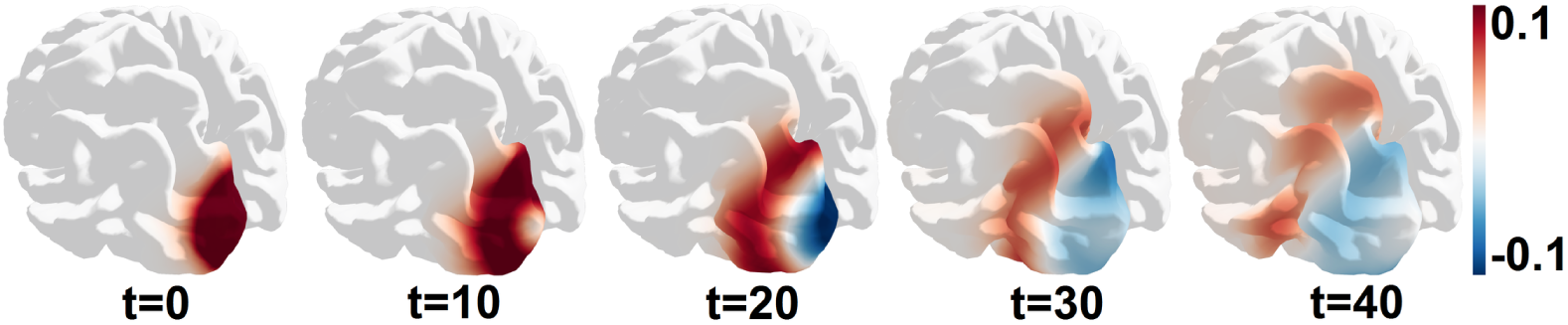
The damped-wave equation on the human connectome gives rise to propagation with characteristic speed and wavelength. Shown are snapshots of simulated cortical activity that is governed by the damped wave equation with time-step *δt* = 1 and parameters *a* = 3 *·* 10^5^, *b* = 5 *·* 10^3^.

**Fig 2.**
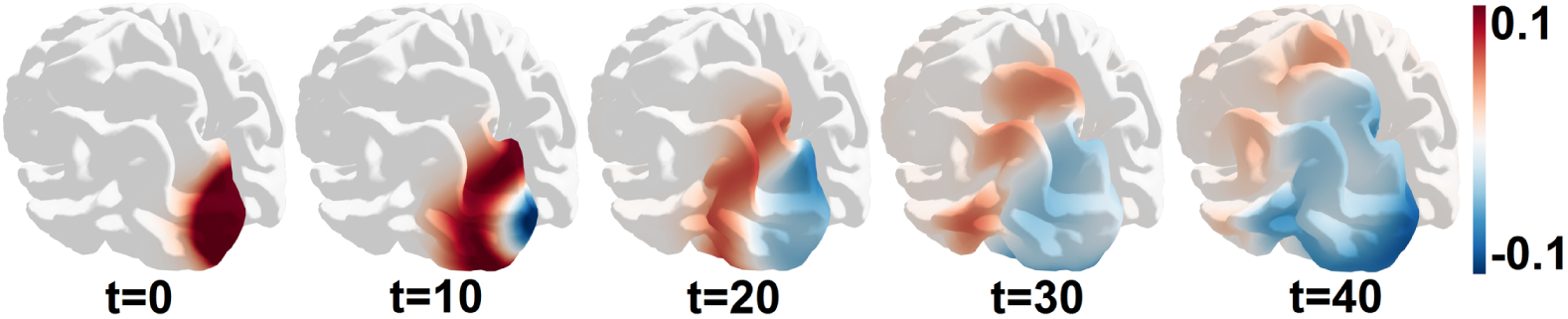
Varying the parameters of the damped-wave equation alters the dynamics of propagation on the human connectome. Shown are snapshots of simulated cortical activity that is governed by the damped wave equation with time-step *δt* = 1 and parameters *a* = 1.5 *·* 10^5^, *b* = 2.5 *·* 10^3^.

**Fig 3.**
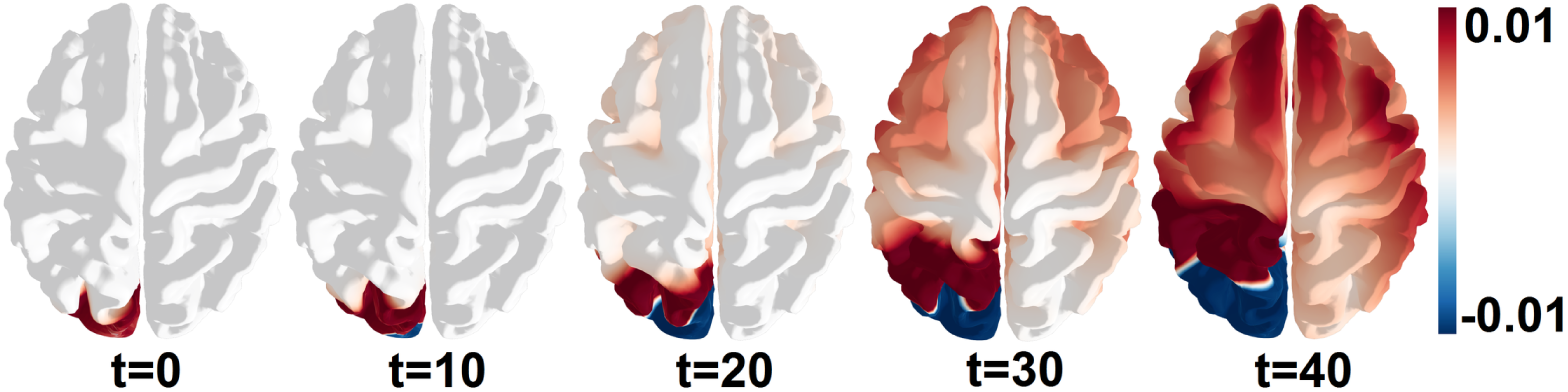
Dynamics of the damped-wave equation on the human connectome include non-local propagation along white-matter fibers. Shown are snapshots of simulated cortical activity that is governed by the damped wave equation with time-step *δt* = 1 and parameters *a* = 1.5 *·* 10^5^, *b* = 2.5 *·* 10^3^.

### Stochastic Wilson-Cowan equations on graphs

The Wilson-Cowan model [20] is a widely used and successful model of cortical dynamics. In this section we show how to use the framework of graph neural fields (*Methods*) to implement the stochastic Wilson-Cowan equations on an arbitrary graphs equipped with a suitable distance metric, and how to compute spatiotemporal observables (CHAOSS). We then illustrate the effects of weighted and non-local graph edges on the dynamics in the simplified context of a one-dimensional graph, before moving on to a real-world application with fMRI data.

#### General formulation

In continuous space, the stochastic Wilson-Cowan model is described by the following system of integrodifferential equations:

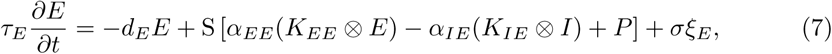

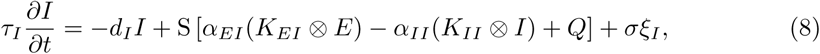

where the symbol ⊗ denotes a convolution integral, and we have omitted for brevity the spatiotemporal dependency of *E*(*x, t*), *I*(*x, t*), *ξ*_*E*_(*x, t*) and *ξ*_*I*_ (*x, t*). In other words, this model posits the existence of two neuronal populations (Excitatory and Inhibitory) at each location in space. The fraction of active neurons in each population (*E*(*x, t*), *I*(*x, t*)) evolve according to a spontaneous decay with rate *d*_*E*_ and *d*_*I*_, a sigmoid-mediated activation term containing the four combinations of population interactions (*E-E, I-E, E-I, I-I*) as well as the external input terms *P* and *Q*, stochastic noise realizations *ξ*_*E*_(*x, t*) and *ξ*_*I*_ (*x, t*) of intensity *σ*, and with the temporal scaling parameters *τ*_*E*_ and *τ*_*I*_. The propagation of activity and interaction among neuronal populations is modeled by spatial convolution integrals with four, potentially different, kernels (*K*_*EE*_, *K*_*IE*_, *K*_*EI*_, *K*_*II*_). For arbitrary symmetric spatial kernels, convolutions can be formulated as linear matrix-vector products on graphs (*Methods* Eq (45)). Therefore, the stochastic Wilson-Cowan equations on graphs reduce to

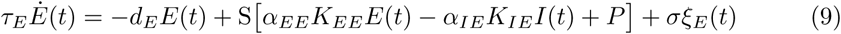

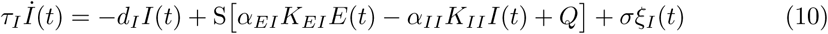

*E*(*t*), *I*(*t*), *ξ*_*E*_(*t*) and *ξ*_*I*_ (*t*) are functions on the graph, i.e. vectors of size *n*, where *n* is the number of vertices in the graph; the convolutions are implemented via the graph-filters *K*_∗∗_, which are matrices of size (*n, n*). In particular, for the case of Gaussian convolutions, the filters are given by (Table 2)

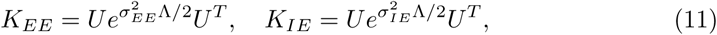

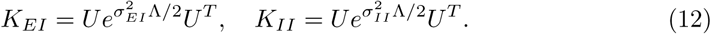

With the distance-weighted graph Laplacian Δ = *U* Λ*U*^*T*^, so that Λ is the diagonal matrix of graph Laplacian eigenvalues, *U* and *U*^*T*^ the corresponding eigenvectors.

This model formulation also allows for non-Gaussian convolution kernels; and the inclusion of a stochastic term allows for characterization of resting-state activity as noise-induced fluctuations about a stable steady-state (*E*^∗^, *I*^∗^) [38].

#### Linear stability analysis

In order to compute meaningful spatiotemporal observables with CHAOSS, it is first necessary to obtain solutions the steady-state equations and their linear stability. For generality and compactness of notation, let us define a new column vector **u**(t) as the concatenation of *E*(*t*) and *I*(*t*); We express **d** for the diagonal matrix containing the damping parameters *d*_*E*_ and *d*_*I*_ and ***τ*** for the matrix containing the timescale parameters *τ*_*E*_ and *τ*_*I*_. The matrix *K* contains the four (arbitrary) graph-filters, **X** is the concatenated vector encoding subcortical inputs *P* and *Q*. We can now write the original system of Eq (9-10) with a single equation

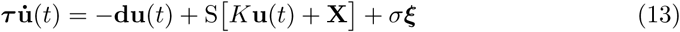

Note that this expression potentially allows for space-dependent model parameters. The steady-state(s) **u**^∗^ can be obtained by setting the time-derivative and noise amplitude *σ* to zero and solving the resulting steady state (matrix) equation:

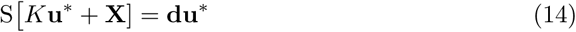

This equation does not have an analytic, exact, closed form solution (in fact, it doesn’t even necessarily have a solution. The sigmoid function is bound between −1 and 1, but **du**^∗^ is not). Furthermore, in the context of a whole-brain model on a mesoscopic connectome, the equation is very high dimensional (twice the number of vertices *n*, with *n* ∼ 18000 in our case). This makes a brute-force numerical approach to the determination of steady states computationally inefficient, especially because it would have to be repeated for each parameter set under examination.

#### Solutions to the steady state equations

If we restrict our analysis to *spatially homogeneous* steady states and *space-independent* model parameters, the steady state equation simplifies to the following 2-dimensional system, rather than the original 2*n*-dimensional Eq (14):

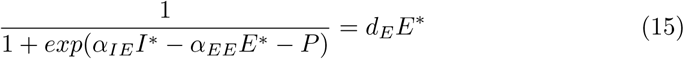

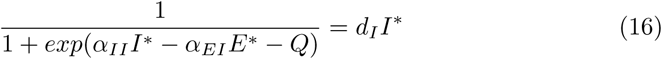

Solutions to this 2-dimensional system can be rapidly obtained numerically for any set of parameters. The biologically valid steady state(s) of the model are given by the solutions (*E*^∗^, *I*^∗^) with *E*^∗^, *I*^∗^ ∈ [0, 1], since *E* and *I* here represent the fraction of active neurons within the respective population. Once a valid steady state is obtained, its stability can be determined through the Jacobian eigenspectrum, and verified with numerical simulations.

#### General Jacobian

To obtain the Jacobian eigenspectrum and determine the stability of a steady state, we have to linearize Eq (13) as

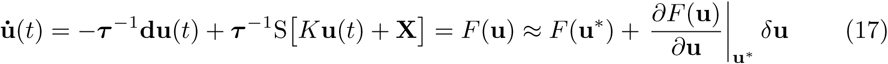

With *F* (**u**^∗^) = 0 by definition since **u**^∗^ is a steady state, and defining a small perturbation abut the steady state *δ***u** = (**u** − **u**^∗^). The Jacobian is then:

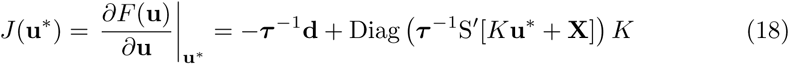

To simplify this expression and allow further analytic progress, we apply the property of the sigmoid derivative S′(*x*) = S(*x*)(1 − S(*x*)); the steady state equation S [*K***u**^∗^ + **X**] = **du**^∗^; and finally denoting with the Hadamard (element-wise) product we obtain

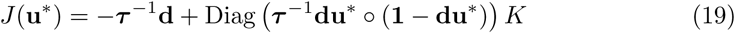

We have thus obtained a general expression for the Jacobian of the Wilson-Cowan model on graphs, which holds also for non-homogeneous steady states and/or space-dependent parameters. In order to evaluate the linear stability of any steady state, it is sufficient plug in the steady state **u**^∗^ = (*E*^∗^, *I*^∗^) solution to Eq (14) in Eq (19), and study the eigenspectrum of the resulting Jacobian.

#### Analytic form of the Jacobian eigenspectrum

On a mesoscopic human connectome graph such as the one we use here, the general Jacobian of Eq (19) is a dense matrix with more than 10^8^ elements. Its eigenspectrum can be calculated numerically, but such a computation is not particularly fast, and has to be repeated for each steady state of each parameter set under examination. However, we restrict the problem to *homogeneous* steady states and *space-independent* model parameters, it is possible to use the properties of the graph Laplacian to obtain an analytic expression for the Jacobian eigenspectrum, which can be computed extremely quickly^2^. These assumptions can be relaxed to allow for eigenmode-dependent^3^ model parameters. Define the scalar, steady-state-dependent parameters:

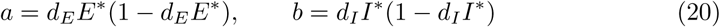

We can use the general Jacobian of Eq (19), with Gaussian kernels, to write explicitly the *linearized* Wilson-Cowan equations for the time-evolution of a perturbation about a homogeneous steady state:

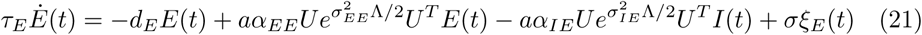

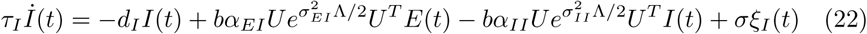

Applying the graph Fourier transform *U*^*T*^, the equations are diagonalized. Each eigenmode of the graph Laplacian therefore behaves independently as a 2-dimensional linear system, with the Jacobian for the *k*^*th*^ eigenmode being

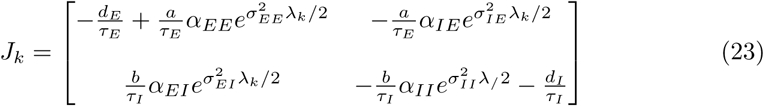

The Jacobian eigenvalues can then be computed directly as

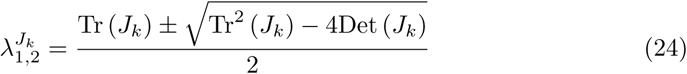

From the Jacobian eigenspectrum thus obtained, we can directly infer the stability character of a steady state. Note that in order to obtain meaningful predictions for spatiotemporal observables, the steady state under examination has to be stable, that is, *J*_*k*_ must have no eigenvalues with positive real parts for all *λ*_*k*_. We have used Gaussian kernels in the derivation, but the result can be straightforwardly generalized to all other kernels. The decoupling of the linearized Wilson-Cowan equations in the graph-Fourier domain has another important consequence: since the dimensionality of the system reduces from *n*^2^ to *n* (where *n* is the number of vertices in the graph), very efficient numerical simulations of the linearized equations can be carried out directly in the graph Fourier domain.

### Wilson-Cowan model CHAOSS

After having obtained steady state solutions, and determined stability character of the steady state, we can apply the *Connectome-Harmonic Analysis Of Spatiotemporal Spectra* (*Methods*) to characterize the spatiotemporal dynamics of resting-state brain activity. CHAOSS predictions, combined with a suitable observation model, can then be compared with empirical neuroimaging data, for example EEG, MEG, or fMRI. Consider linearized stochastic graph neural field equations in the graph Fourier domain:

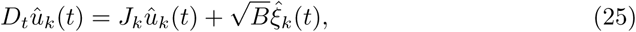

Where *k* = 1, …, *n* indexes the graph Laplacian eigenmodes. For Wilson-Cowan model (Eq (21-22)), we have: *D*_*t*_ = *d/dt, J*_*k*_ is given in Eq (23) for the case with Gaussian kernels, and *B* is

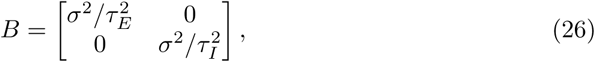

In terms of the elements of the matrices *J*_*k*_ and *B*, the theoretical prediction for the harmonic-temporal power spectrum of the excitatory neural population activity becomes (*Methods* Eq (72)):

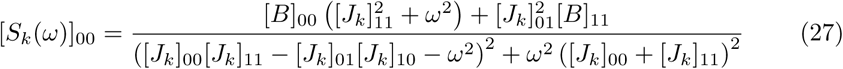

The double-digits numerical subscripts in Eq (27) refer to the row-column element of the respective matrix. Eq (27) can be used to compute the separate harmonic and temporal power spectra of the model, as well as the correlation and functional connectivity matrices (*Methods* Eq (77)). Equivalent formulas for the inhibitory population can also be similarly derived.

By integrating [*S*_*k*_(*ω*)]_00_ over all temporal frequencies, an explicit expression for the harmonic power spectrum of excitatory activity can be obtained:

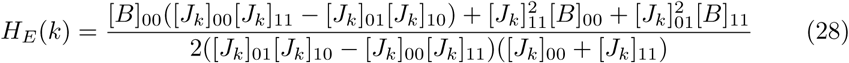

Eq (27) and (28) represent a general result that does not only apply to the Wilson-Cowan model. In fact, these equations describe the harmonic-temporal and harmonic power spectra of stochastic equilibrium fluctuations for the first population of any graph neural field model with two interacting populations and a first-order temporal differential operator.

#### Effects of distance-weighting and long-range connectivity

We characterize the effects internode spacing and the presence of long-range connectivity on model dynamics by implementing the Wilson-Cowan model on a one-dimensional graph with 1000 nodes. Numerical simulations were carried out with a time-step *δt* = 5 *·* 10^−5^*s* and an observation time of 10^5^ time-steps, which corresponds to 5*s* of simulated activity.

To show the effects of distance-weighting in graph neural fields, we note how, for the parameter set of S1 Table, increasing the distance between nodes leads to the emergence of an oscillatory resonance that eventually destabilizes and gives way to limit-cycle activity. Keeping the number of nodes constant, increasing the internode spacing *h* alters the stability of the steady state from broadband activity (*h* = 10^−5^m), to oscillatory resonance (*h* = 10^−4^m), to oscillatory instability (*h* = 2*·*10^−4^m). The spatial and temporal power spectra for the case *h* = 10^−4^m are shown Figure 4A and B, respectively. These results demonstrate that by using the weighted graph Laplacian, the dynamics of graph neural fields becomes dependent on how metric properties of the graph i.e. on how the graph is embedded in three-dimensional space. If the combinatorial graph Laplacian is used, as done in [9], the dynamics only depends on the topology of the cortical mesh and in this sense does not take into account the physical properties of the cortex.

**Fig 4.**
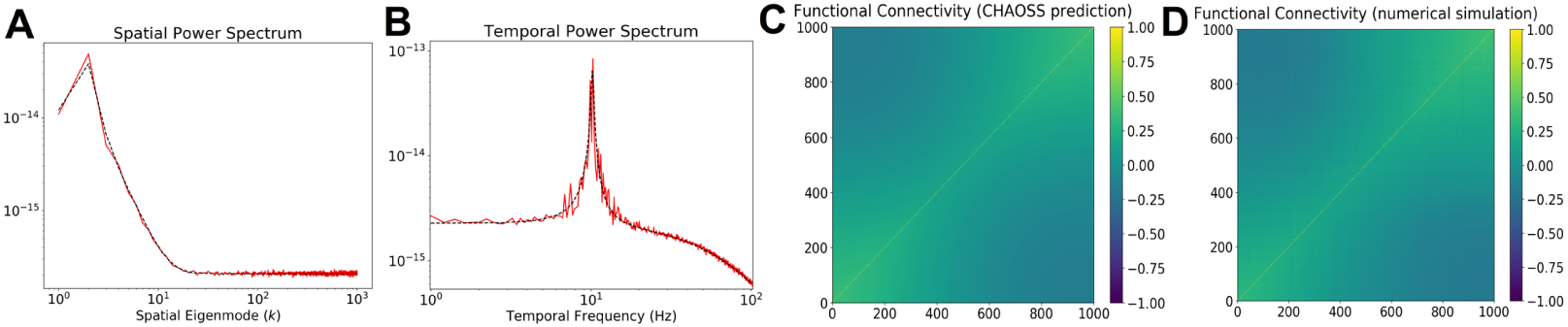
Effects of internode spacing on graph neural field dynamics. Panels A and B show, respectively, the spatial and temporal power spectrum of equilibrium fluctuations in the one-dimensional graph for internode spacing *h* = 10^−4^*m*. The dotted black correspond to the theoretical prediction and the red lines are obtained through numerical simulations. Panels C and D show the model functional connectivity as obtained, respectively, by analytic predictions and numerical simulations.

The presence of *fast*, long-range connectivity can impact the power spectrum and functional interactions of equilibrium fluctuations, as well as the stability of steady-states. To illustrate this, we add a single non-local edge between nodes 250 and 750 to the one-dimensional graph with *h* = 10^−4^m. The Euclidean distance between these two nodes is 500 *h* = 5*·*10^−2^m= 5cm. In the healthy brain, myelination of white-matter fibers allows long-range (cortico-cortical) activity propagation to take place at speeds ∼ 200 times greater than local (intra-cortical) propagation [21]. To model myelination, we set the length of the non-local edge to be the Euclidean distance between the nodes, divided by a factor of 200 (similarly to the construction of the human connectome graph Laplacian, where the length of cortico-cortical edges is set to be their path-length distance along DTI fibers, divided by a factor of 200). Therefore, the effective length of the non-local edge is 2.5*·*10^−4^m. Figure 5 shows the effects of the presence of the non-local edge on the spatial and temporal power spectra of the equilibrium fluctuations.

**Fig 5.**
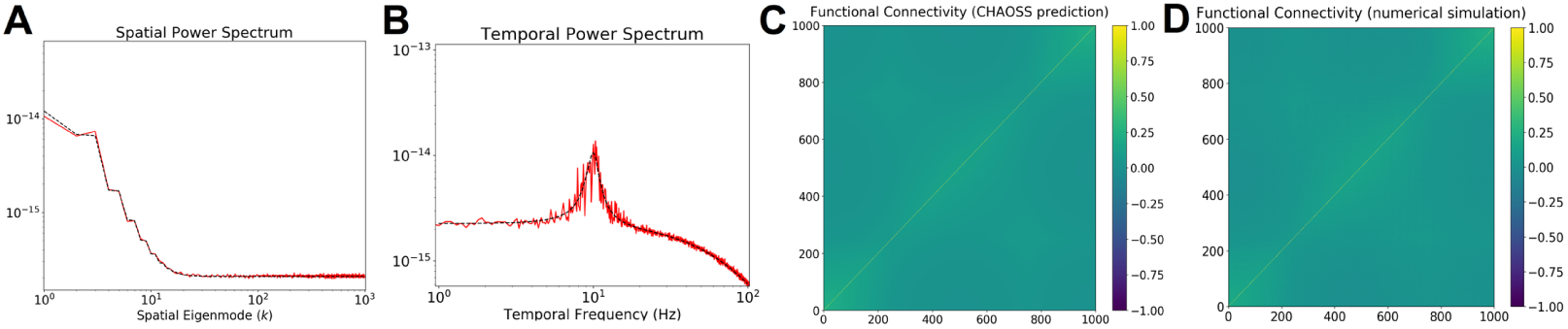
Suppression of oscillatory resonance by long-range connectivity. Panels A and B show, respectively, the spatial and temporal power spectrum of equilibrium fluctuations in the one-dimensional graph for internode spacing *h* = 10^−4^*m* after the addition of a non-local edge between nodes 250 and 750. Compare with Figure 4 to note the visible suppression of oscillatory resonance, and the slight change in functional connectivity engendered by a single non-local edge. The dotted black correspond to the theoretical prediction and the red lines are obtained through numerical simulations. Panels C and D show the model functional connectivity as obtained, respectively, by analytic predictions and numerical simulations.

The most pronounced effect is damping of the oscillatory resonance in the temporal power spectrum, thus rendering the fluctuations more stable. Furthermore, the edge leads to a faint but discernible alteration in the functional connectivity (see Panels C and D).

Interestingly, when the model operates in the pathological i.e. non-stable regime (*h* = 2*·*10^−4^m), addition of a single non-local edge stabilizes the steady state, thus leading to healthy equilibrium fluctuations (see Figure 6). The non-local edge also creates a visible increase in the functional connectivity between the nodes involved, and a change in the pattern in neighboring nodes (Panels C and D). As noted above, these significant effects of long-range connectivity are observed if the effective length of the non-local edge is small enough for non-local activity propagation to interact with local activity propagation. For these one-dimensional simulations, this happens if the speed factor is larger than ∼50.

**Fig 6.**
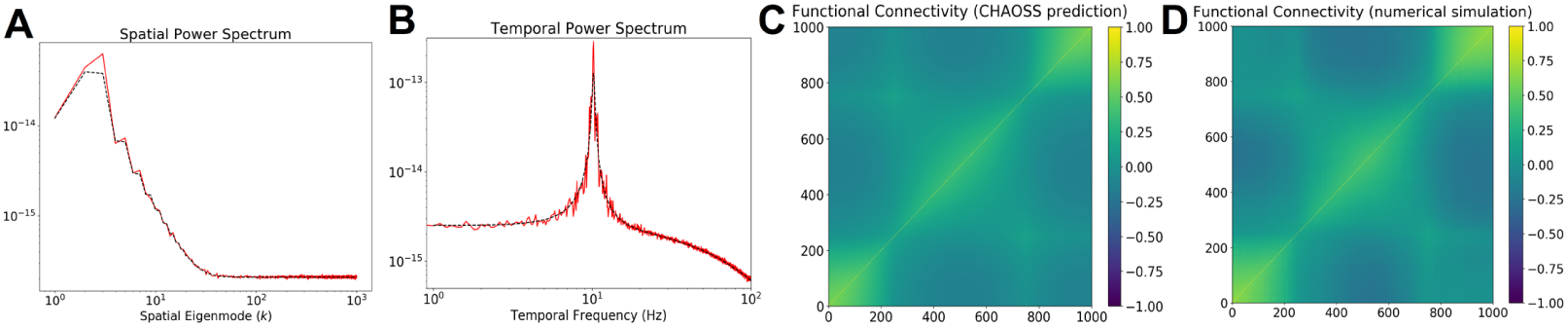
Abortion of pathological oscillations by long-range connectivity. Panels A and B show, respectively, the spatial and temporal power spectrum of equilibrium fluctuations in the one-dimensional graph for internode spacing *h* = 2*·*10^−4^*m* after the addition of a non-local edge between nodes 250 and 750. Before the addition, the dynamics was placed in an unstable limit-cycle regime. The dotted black correspond to the theoretical prediction and the red lines are obtained through numerical simulations.Panels C and D show the model functional connectivity as obtained, respectively, by analytic predictions and numerical simulations.

### Application to resting-state functional MRI

To illustrate a real-world application of graph neural fields, we implement the stochastic Wilson-Cowan model on the connectome graph of a single subject, and use it reproduce the harmonic power spectrum observed in resting-state fMRI data. The harmonic power spectrum of resting-state fMRI was computed, according to its definition, as the temporal mean of the squared graph-Fourier transform of the fMRI timecourses. To regularize the empirical fMRI harmonic power spectrum, we computed its log-log binned median with 300 bins, following [22]. Eigenmodes above *k* ∼ 15000 contained artifacts due to reaching the limits of fMRI spatial resolution, and were thus removed. The model power spectra were binned in the same way. The parameters of the Wilson-Cowan model were optimized with a basinhopping procedure, aiming to minimize the residual difference between observed and predicted spectra. In the fitting, we allowed for a linear rescaling of the theoretical spectrum so that

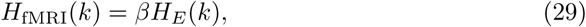

where *H*_fMRI_(*k*) is the harmonic power spectrum of the fMRI data, *H*_*E*_(*k*) is the harmonic power spectrum of the excitatory population in the Wilson-Cowan model, and *β* is a rescaling parameter. Fig 7 shows the best fitting theoretical harmonic power spectra, together with the empirical fMRI spectrum. The model is able to reproduce the experimental spectrum. The parameters values obtained from the fitting are reported in S2 Table and appear consistent.

**Fig 7.**
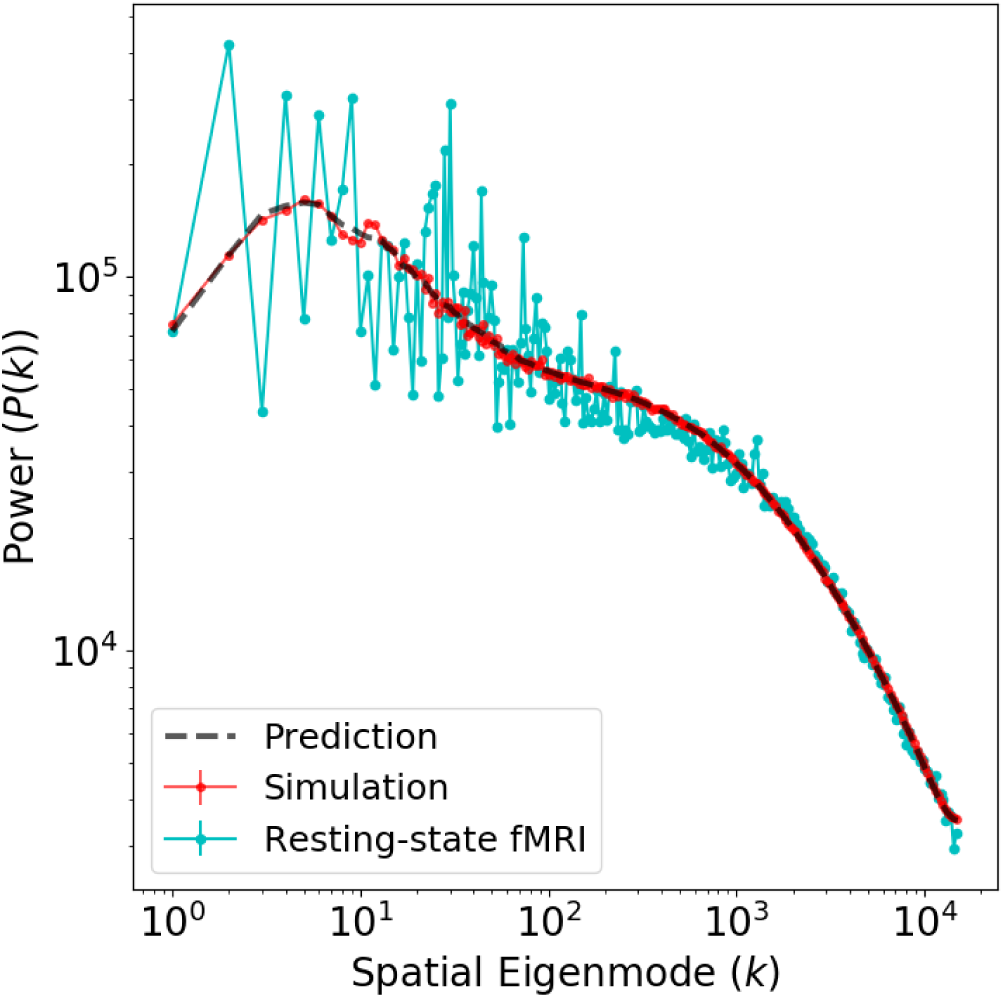
Stochastic Wilson-Cowan graph neural field model predicts the experimental harmonic power spectrum. Shown is the empirically observed BOLD-fMRI harmonic spectrum (cyan line), together with the theoretical (dotted black line) and numerical (red line) predictions from the stochastic Wilson-Cowan equations. The numerical prediction was obtained by taking the median of three independent simulations.

To verify the accuracy of predictions obtained with the connectome-harmonic analysis of spatiotemporal spectra, we implement numerical simulations of the linearized Wilson-Cowan equations on the human connectome graph. Simulations were carried out with a time-step value *δt* = 10^−4^*s* and an observation time of 10^6^ time-steps, corresponding to 100*s* of simulated brain activity. Figure 8 shows snapshots of the simulated resting-state activity at several different times. Note that the two hemispheric meshes appear physically separate, but inter-hemispheric propagation is allowed through white-matter callosal tracts (not shown in the figure).

**Fig 8.**
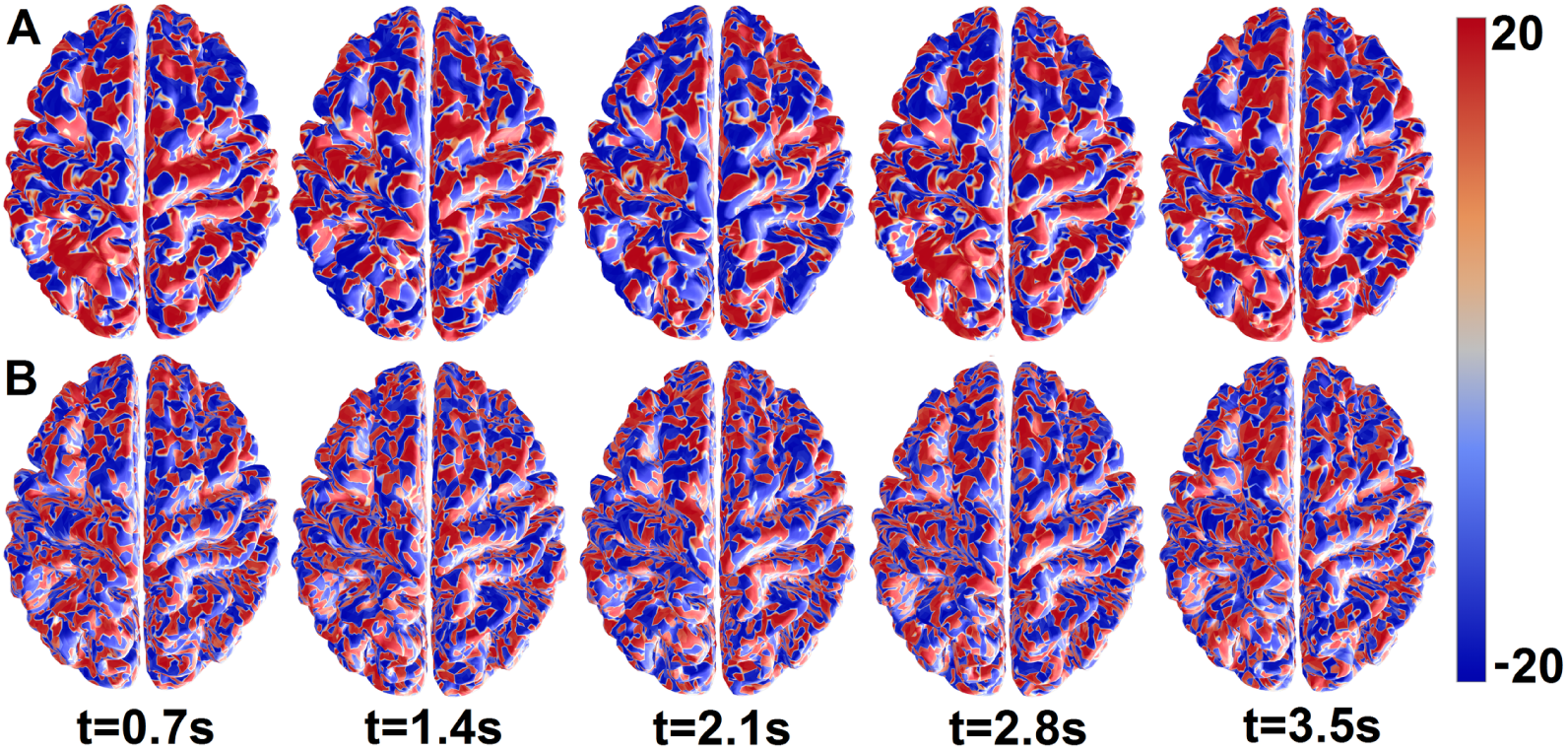
Measured and simulated resting-state brain activity. Panel A shows resting-state brain activity, as fluctuations of the BOLD fMRI signal about the mean at each node. Panel B shows snapshots of activity from the stochastic Wilson-Cowan equations simulated using the parameters of Table in S2 Table. The model activity was temporally downsampled to match the TR of fMRI data, and rescaled by 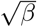 to match the scale of the BOLD signal. No spatial or temporal smoothings were applied.

## Methods

### Laplacian operators on graphs

In this section we provide a derivation of the weighted graph Laplacian operator in terms of graph differential operators. The weighted graph Laplacian is distinguished from the combinatorial graph Laplacian often used in analysis studies [9], as it allows physical properties of the cortex to be taken into account, which is necessary to implement “realistic” graph neural field models.

#### The combinatorial Laplacian

Consider an undirected graph with *n* vertices. The graph’s binary *adjacency matrix Ã* is defined by

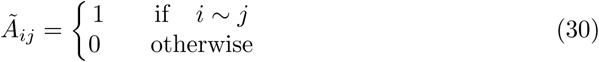

where *i* ∼ *j* means that vertices *i* and *j* are connected by an edge. The graph’s *degree matrix* 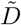 is a diagonal matrix whose diagonal entries are defined by

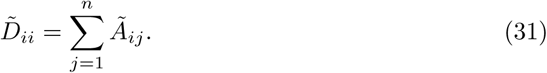

It hence counts the number of edges for each vertex *i*. The binary or *combinatorial graph Laplacian*, denoted by 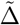, is defined as

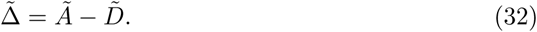

The combinatorial graph Laplacian and its normalized version do not carry information about the distances between cortical vertices and therefore are invariant under topological but non-isometric deformations of graph. Neural activity modeled in terms of the combinatorial graph Laplacian therefore is a topological graph invariant, whereas real neural activity does depend on the metric properties of the graph. The combinatorial graph Laplacian, however, can be adjusted so as to take into account the metric properties of the graph, yielding the *weighted graph Laplacian*. Below, we provide a derivation of the weighted graph Laplacian in terms of the graph directional derivaties of a graph function.

#### The weighted Laplacian

Let *f* be a *graph function* i.e. a function defined on the vertices of a graph and let *M* be the graph’s distance matrix. Thus, the (*i, j*)^*th*^ entry *M*_*ij*_ of *M* equals the distance between nodes *i* and *j* in a particular metric. We note that for this derivation, it is irrelevant how *M* is obtained. In the context of connectomes, the elements of *M* can be defined in terms of suitably scaled Euclidean distances, geodesic distances over the cortical manifold, or as the lengths of the white matter fibers connecting the vertices. The *first-order graph directional derivative* ∂_*j*_*f*_*i*_ of *f* at vertex *i* in the direction of vertex *j* is defined as

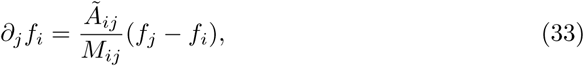

Note that according to this definition, ∂_*j*_*f*_*i*_ = 0 if vertex *j* is not connected to vertex *i*, and that ∂_*i*_*f*_*i*_ = 0. Also note that ∂_*j*_ is a linear operator on the vector space of graph signals. Furthermore, since 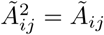, the *second-order graph directional derivative* 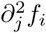 of *f* at vertex *i* in the direction of vertex *j* is defined as

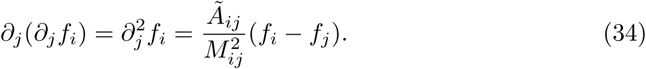

Following the definition of the Laplacian operator in Euclidean space as the sum of second-order partial derivatives, the *weighted graph Laplacian*, denoted as Δ is defined as ^4^

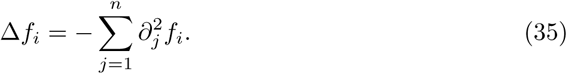

To see the relation with the combinatorical graph Laplacian, we note that Δ can be written in matrix form as

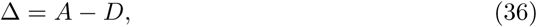

where *A* and *D* are the *weighted adjacency matrix* and *weighted degree matrix*, respectively, which are defined as 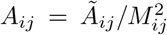 and 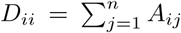, respectively. Thus, the weighted graph Laplacian can be obtained by using the weighted versions of the adjacency and degree matrices in the definition of the combinatorical graph Laplacian.

### Convolution of graph signals

In order to define graph neural fields, we need to have a graph-theoretical analog of the continuous spatiotemporal convolution (*K* ⊗ *u*)(*x, t*)

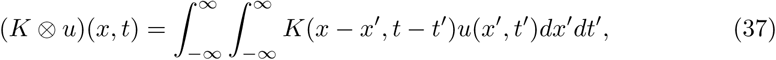

To obtain this, we use the convolution theorem to represent the convolution in the spatiotemporal Fourier domain as 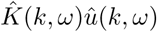, where *k* denotes wavenumber amd *ω* denotes temporal frequency. When the kernel is real-valued and spatially symmetric, its Fourier transform is real-valued and even in *k*, so that 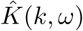 can be viewed as a function of − *k*^2^. To generalize the convolution operator to graphs, we use the fact that −*k*^2^ is the eigenvalue of the spatial Fourier basis function *e*^*ikx*^ under the Laplace operator *d*^2^*/dx*^2^. Since the Laplace operator on a graph is given by its Laplacian matrix Δ, the graph filter corresponding to the spatiotemporal convolution (*K* ⊗ *u*)(*x, t*) can be defined by substituting the eigenvalues of Δ for the values −*k*^2^ in 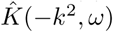.

#### The graph Fourier transform

We consider a graph with *n* vertices and Laplacian matrix Δ, together with an eigendecomposition Δ = *U* Λ*U*^*T*^, where *U* is an *n × n* orthogonal matrix containing an eigenbasis of Δ in its columns and Λ is the *n × n* diagonal matrix that contains the corresponding eigenvalues *λ*_1_ ≥ *λ*_2_ ≥, *…*, ≥ *λ*_*n*_ ≥ 0 on its diagonal. The *graph Fourier transform* of a function *u*(*t*) on the graph is now defined by

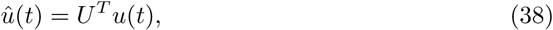

where the transformation *U*^*T*^ expresses *u*(*t*) in the eigenbasis of Δ. The vertex-domain signal *u*(*t*) can be recovered again by applying the *inverse graph Fourier transform U* ^−*T*^ = *U* to *û*(*t*). For clarity, note that the graph Fourier transform is not related to the temporal Fourier transform and that *u*(*t*) does not have to depend on time to apply it. In particular, for grid graphs (i.e. graphs whose drawing, embedded in some Euclidean space, forms a regular tiling), the graph Fourier transform is equivalent, in the continuum limit, to the spatial Fourier transform in Euclidean space. However, the graph Fourier transform as defined here is a more general concept, since it can be applied to arbitrary nondirected graphs, possibly with non-local edges, such as the human connectome.

#### Filtering of graph signals

Let *K*(*x, t*) be a kernel with Fourier transform 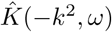 and let *u*(*t*) be a graph signal with spatial Fourier transform *û*(*t*). We denote the temporal Fourier transforms of *u*(*t*) and *û*(*t*) by *u*(*ω*) and *û*(*ω*), respectively. We define the graph kernel 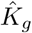 associated with the continuous kernel *K*

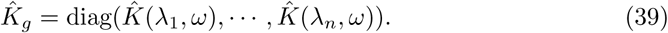

In the graph Fourier and temporal frequency domains, the filtered signal is hence per definition given by

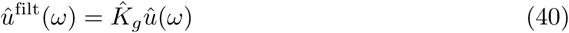

Applying the inverse graph Fourier transform *U*, we obtain the filtered signal in the graph domain

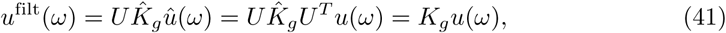

where we have defined 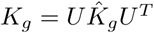, the graph domain representation of the filter. In other words, *K*_*g*_ is the graph filtering operator corresponding to the continuous kernel *K*.

The above representations of the kernel and the filtered signal are in the temporal frequency domain. To obtain the corresponding time domain representations, we note that the *i*^*th*^ entry of *u*^filt^(*ω*) is given by

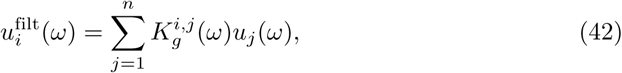

where 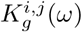 denotes the (*i, j*)^*th*^ entry of *K*_*g*_(*ω*). Using the convolution theorem, the time domain representation of the filtered signal is

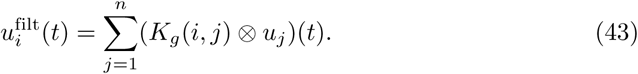

Collecting the terms for all *n* entries in a column vector yields the filtered signal in the graph and temporal domain and we write it as (*K*_*g*_ ⊗_*g*_ *u*)(*t*):

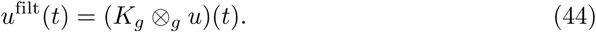

In case of a purely spatial kernel *K*(*x, t*) = *K*(*x*), with *K*(*x*) symmetric and with spatial Fourier transform 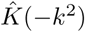, the graph kernel 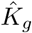 is independent of frequency so that the filtering operator reduces to the frequency-independent linear transformation

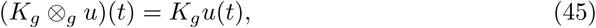

In case of a purely temporal kernel *K*(*x, t*) = *g*_Θ_(*t*), with *g*_Θ_(*t*) = *g*(*t*)Θ(*t*), where *g*(*t*) is the temporal kernel, and Θ(*t*) the Heaviside step function, which ensures that integration is temporally causal. Graph filtering reduces to a temporal convolution:

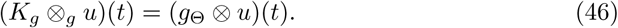

Lastly, in case of a separable kernel *K*(*x, t*) = *w*(*x*)*g*_Θ_(*t*), the graph kernel decomposes as

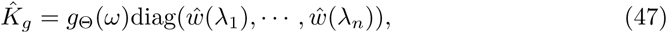

so that the filtered signal is given by

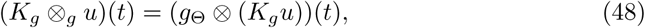

In this case, the filtering occurs in two stages: the signal of first spatially filtered by *K*_*g*_ and subsequently temporally filtered by the convolution with *g*_Θ_(*t*).

#### Examples of graph kernels

We have defined spatiotemporal graph kernels for maximal generality, but in practice, spatial kernels are particularly efficient and simple to use, since they reduce to simple linear products in the graph domain. Table 2 lists several commonly used continuous spatial kernels and their equivalent on graphs. The graph Laplacian converges to the Laplacian operator in the continuum limit; therefore on grid graphs (i.e. graphs whose drawing, embedded in some Euclidean space, forms a regular tiling), the graph filters act simply like discretized versions of their continuous counterparts. For example, applying the graph Gaussian kernel to a function defined on a 2-dimensional square-grid graph is equivalent to blurring/smoothing a 2-dimensional image with a spatial Gaussian kernel of the same size, and closed boundary conditions (open boundaries can be implemented by extending the graph beyond the image size, and periodic boundaries by adding edges connecting nodes on opposite sides). More relevantly to the current work, this approach generalizes to arbitrary non-grid graphs, potentially with non-local edges, such as the human connectome, and is therefore more general than grid-based discretizations of continuous convolution kernels.

#### Reaction-diffusion neural activity models

In this section we show how graph filters can also be used to implement the graph equivalents of neural activity models that can be directly written as partial differential equations [18, 23] and, among others, comprise damped wave and reaction-diffusion equations.

Consider the following scalar reaction-diffusion model

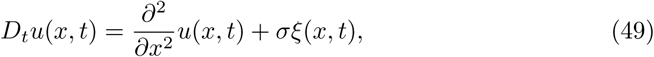

where *D*_*t*_ denotes a temporal differential operator that describes local reactions, ∂^2^*/*∂*x*^2^ is the diffusion term and *σξ*(*x, t*) is an external forcing function. To obtain the corresponding graph equation we first transform Eq (49) to the spatiotemporal Fourier domain:

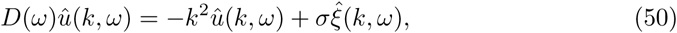

where *D*(*ω*) denotes the Fourier transform of *D*(*t*), and subsequently solve for *û*(*k, ω*):

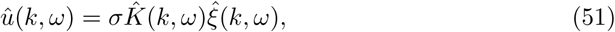

where the spatiotemporal kernel 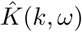 is given by

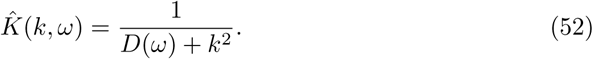

The corresponding graph kernel is given by

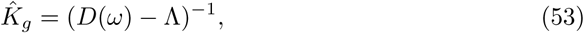

where Λ denotes the diagonal matrix containing the eigenvalues of the weighted graph Laplacian Δ. Applying 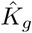 to the input gives

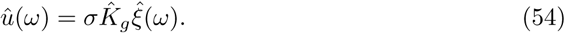

Transforming back to the graph-temporal domain to obtain *u*(*t*) = *σ*(*K*_*g*_ ⊗ *ξ*)(*t*). To obtain the full system of differential equations, we note that

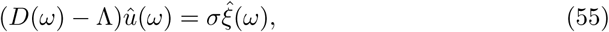

and transform this equation back to the spatial domain to obtain the following system of ordinary differential equations:

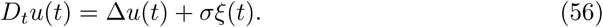

This shows that a reaction diffusion equation can directly be defined on a graph by replacing the Laplace operator in continuous space by the weighted graph Laplacian Δ, and solved by computing a suitable graph filter.

### Graph neural fields

#### Continuous neural fields

Neural field models describe the dynamics of cortical activity *u*(*x, t*) at time *t* and cortical location *x* ∈ Ω. Here, Ω ∈ ℝ^3^ denotes the cortical manifold embedded in three-dimensional Euclidean space. Depending on the physical interpretation of the state variable *u*(*x, t*), neural fields come in two types, which we will refer in the rest of the text as *Type 1* and *Type 2*. This short description is by no means meant to be exhaustive, and only contains the required background to define graph neural fields; comprehensive treatments of continuous neural fields are provided in [18, 23].

In Type 1 neural fields [18], the state variable *u*(*x, t*) describes the average membrane potential at location *x* and time *t*. The general form of a neural field model of Type 1 is

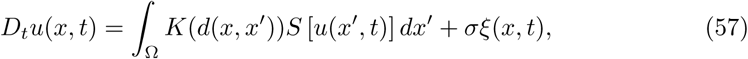

where *σξ*(*x, t*) is the external input, *d*(*x, x*′) is the geodesic distance between cortical locations *x* and *x*′, *K* is the spatial kernel of the neural field that describes how the firingrate *S*[*u*(*x*′, *t*)] at location *x*′ affects the voltage at location *x*, and *S* is the firing-rate function that converts voltages to firing-rates. Furthermore, *D*_*t*_ is a temporal differential operator that models the synaptic dynamics. In modelling ongoing cortical activity, *ξ*(*t, x*) is usually taken to be a stationary stochastic process. Following earlier studies, we will assume *ξ*(*t, x*) to be spatiotemporally white-noise, i.e.

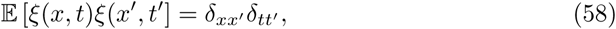

where 𝔼 denotes expectation value and δ_*aa′*_ is the Dirac delta function, which is defined by 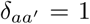 if *a* = *a*′ and zero otherwise. Note that in principle, colored noise could be used as well. The distance function *d*(*x, x*′) between cortical locations *x* and *x*′ as well as the integration over the cortical manifold Ω assume that Ω is equipped with a Riemannian metric. A natural metric is the Euclidean metric induced by the embedding of the cortical manifold in three-dimensional Euclidean space.

In Type 2 neural field models [24, 25], the state variable *u*(*x, t*) denotes the fraction of active cells in a local cell population at location *x* and time *t* and hence takes values in the interval [0, 1]. Type 2 neural field models have the form

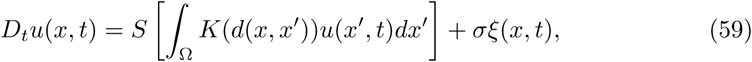

where *S* denotes the activation function that maps fractions to fractions and hence takes values in the interval [0, 1] and thus has a different interpretation than the firing-rate function in Type 1 neural field models. Note that the only difference between Type 1 and Type 2 neural field models is the placement of the non-linear function *S*. Most neural field models used in practice are obtained by coupling two or more field models, where each model corresponds to a different neural population. For example, the state variable of the Wilson-Cowan neural field model is two-dimensional and its components correspond to excitatory and inhibitory neural populations. In the rest of the text, we will assume that *K* is normalized to have unit surface area.

In theoretical studies on neural field models, the cortex is usually assumed to be flat:, i.e. Ω = ℝ^2^ (cortical sheet) or Ω = ℝ^1^ (cortical line) or a closed subset thereof.

The major simplification that occurs in this case is that the cortical metric reduces to the Euclidean metric:

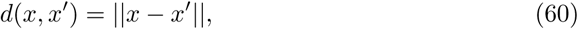

and, as a consequence, the integrals in Eq (57) and (59) reduce to convolutions so that Fourier methods can be used in the models’ analysis (however, see [27] for a detailed theoretical study of a neural field model on the sphere). Note that in Euclidean space, the kernel *K* is symmetric, i.e. *K*(− *x*) = *K*(*x*) for all *x* ∈ ℝ. The definition of graph neural fields as given in Section *Graph neural fields* relies on this fact as well. For ease of notation, below we set Ω = ℝ^1^ so that *d*(*x, x*′) = |*x* − *x*′|. Note that the double integrals in Eq (57) and (59) can be written as convolutions, and hence it will be possible to implement them on graphs using the previously derived definitions of graph spatiotemporal convolutions.

#### Graph neural fields

Neural fields given by Eq (57) and (59) can be defined on an arbitrary graph by replacing the spatial filter *K* by its graph-equivalent *K*_*g*_ as defined in Section *Convolution of graph signals*. Thus, a *graph neural field of Type 1* is a model of the form

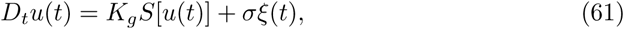

and

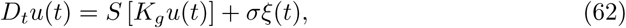

where *u*(*t*) and *ξ*(*t*) are graph functions. Thus, the graph-theoretical analogs of the continuous neural fields given by Eq (57) and (59) are ordinary differential equations. When more than one type of neural population is included or when the differential operator *D*_*t*_ is of order higher than one, the continuous neural fields reduce to *systems* of ordinary differential equations.

The continuous neural fields in Eq (57) and (59) are described by partial integro-differential equations in which the integration in done over space. They can also be described by spatiotemporal integral equations by viewing the differential operator as a temporal integral, which leads to a more general class of continuous neural fields. Using the spatiotemporal convolution on graphs, this more general class of neural fields can be formulated on graphs leading to a systems of temporal integral equations. To make this explicit, we use the definition of the spatiotemporal graph filtering operator *K*_*g*_ ⊗ to write out the *i*^*th*^ component of *u*_*i*_:

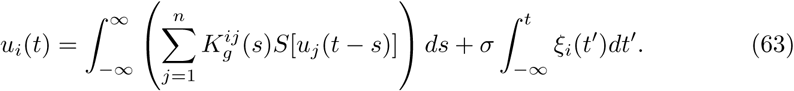

Thus, the spatiotemporal integrals in continuous neural fields are replaced by temporal integrals in graph neural fields and the spatial structure of the continuous neural field is incorporated into the graph kernels 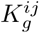. The same applies to neural fields of Type 2. Furthermore, for continuous neural fields of Type 1 with separable kernels, and for special choices of the temporal component of the kernel, the spatiotemporal integral equation can be reduced to a partial integro-differential equation [23, 26]. For graph neural fields there exists an equivalent subset of models that can be represented by a system of ordinary integro-differential equations.

In case of a purely spatial kernel *K*(*x, t*) = *w*(*x*) we obtain the following systems of ordinary differential equations:

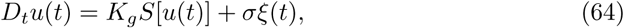

and

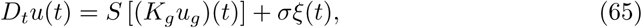

respectively, and in case of a purely temporal kernel *K*(*x, t*) = *g*_Θ_(*x*) we obtain the following systems of ordinary differential equations:

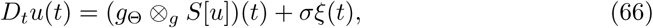

and

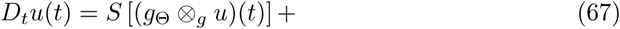

respectively. In case of a separable kernel *K*(*x, t*) = *w*(*x*)*g*_Θ_(*t*) we obtain the following systems of ordinary differential equations:

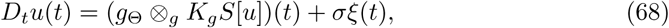

and

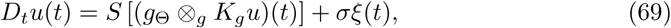

respectively.

### Relating graph neural fields to experimental observables

#### Connectome-harmonic analysis of spatiotemporal spectra (CHAOSS)

To characterize the spatiotemporal statistics of resting-state brain dynamics, we derive analytic predictions for harmonic and temporal spectra and functional connectivity matrices. The derivation relies on the fact that, for space-independent parameters, linearized graph neural field equations decouple in the graph Fourier domain^5^. Each of the *n* graph Laplacian eigenmodes behaves independently like the following *N* -dimensional linear system, where *N* is the number of neuronal population types (*N* = 2 for the Wilson-Cowan model):

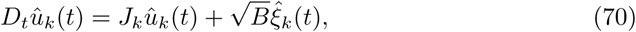

where *ξ*_*k*_(*t*) is an *N* -dimensional uncorrelated white noise, *B* is an *N* -dimensional diagonal matrix containing the noise intensity of the *N* neural populations, *D*_*t*_ is a temporal differential operator, and *J*_*k*_ is the Jacobian of the *k*^*th*^ eigenmode. Taking the temporal Fourier transform we obtain

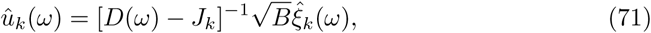

where *D*(*ω*) denotes the temporal Fourier transform of *D*_*t*_. Abbreviating the graph filter 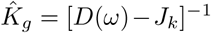, the cross-spectral matrix *S*_*k*_(*ω*) of the *k*^*th*^ eigenmode is given by:

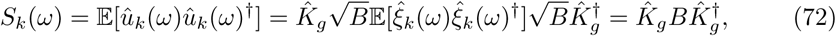

where, *†* denotes the conjugate transpose and E denotes the expected value. Colored noise can be modeled by letting *B* depend on *ω*, although this is usually not done in neural field modelling studies. Another possible generalization is to let *B* depend on the eigenmode *k*.

Eq (72) gives a closed-form expression for the *N* -dimensional cross-spectral matrix of the *k*^*th*^ eigenmode, where *N* is the number of neuronal populations in the model. Hence, its diagonal entries [*S*_*k*_(*ω*)]_*pp*_, *p* = 0, …, *N* − 1 describe the power of the *p*^*th*^ neuronal population in the *k*^*th*^ eigenmode, at temporal frequency *ω*. The temporal power spectrum *T*_*p*_(*ω*) of the *p*^*th*^ neuronal population is obtained by summing over eigenmodes:

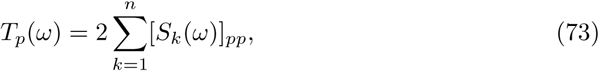

Where the factor of 2 arises because on graphs, *k* ranges only over positive integers between 1 and *n* (*n* is the number of eigenmodes). Similarly, the harmonic power spectrum of the *p*^*th*^ neuronal population *H*_*p*_(*k*) is obtained by integrating over the temporal frequency *ω*:

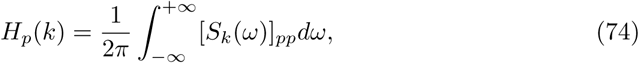

where 1*/*2*π* is a normalization constant. When combined with a suitable observation model, these predictions can be compared with or fitted to experimental data from different neuroimaging modalities.

#### Functional connectivity

Furthermore, it is possible to compute the correlation matrix of brain activity for each neuronal population. To construct the covariance matrix of a neuronal population activity Σ_*p*_ across all graph vertices, we first construct the covariance matrix 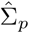 in the graph Fourier domain. Because of the independence of eigenmodes, the covariance matrix of the *p*^*th*^ population (at lag zero) in the graph Fourier domain 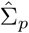 is a diagonal matrix with the elements along the diagonal being the values of the spatial power spectrum.

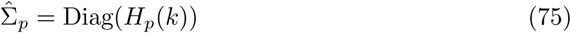

The covariance matrix across all vertices is obtained by transforming back to the vertex domain:

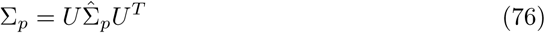

The functional connectivity (correlation) matrix *F*_*p*_, which is often used in fMRI restingstate studies, is obtained by normalizing the covariance matrix

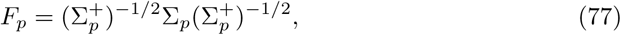

where 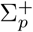 denotes Σ_*p*_ with all off-diagonal entries set to zero. Seed-based connectivity of the *j*^*th*^ node is measured by the *j*^*th*^ row (or column) of *F*_*p*_.

#### Coherence matrix

From the linearized model equations one can also derive the coherence matrix, which measures the strength and latency of linear interactions between pairs of vertices as a function of frequency *ω* and is often used in EEG and MEG studies [28]. If the noise is assumed to be white, non-linear connectivity measures such as the phase-locking value and amplitude correlations can be analytically computed from the coherence matrix [29]. For simplicity we derive the coherence matrix for the special case of a single population and note that the generalization to multiple populations is straightforward.

The derivation of the coherence matrix is similar to that of the functional connectivity, and starts by expressing the linearized model equations in the vertex domain:

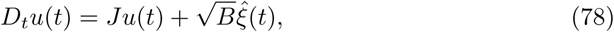

where the *n*-dimensional matrix *J* denotes the Jacobian matrix in the vertex-domain. Transforming Eq. (78) to the temporal Fourier domain and taking expectations yields the cross-spectral matrix *S*_*v*_(*ω*) in the vertex domain:

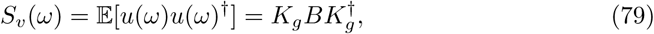

where *K*_*g*_ = [*D*(*ω*) − *J*]^−1^. The coherence matrix *C*(*ω*) is obtained by normalization of the cross-spectral matrix in the vertex domain:

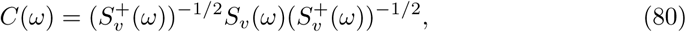

where 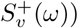 denotes *S*_*v*_(*ω*) with its off-diagonal entries to zero. The (*i, j*)^*th*^ entry of *C*(*ω*) is the coherence between the cortical activity at vertex *i* and *j*.

### Anatomical and functional data

We use the same structural MRI and DTI data as in [9], obtained from the Human Connectome Project (https://db.humanconnectome.org/) to construct the anatomical connectome graph of a single subject. In short, MRI data is employed to obtain intra-cortical graph edges based on the surface mesh; DTI data is employed to add long-range cortico-cortical edges to the graph. The main difference with [9] is that instead of constructing the combinatorial (binary) graph Laplacian, here we construct a distance-weighted graph Laplacian. This allows us to take into account physical distance properties of the cortex that are relevant for graph neural fields, and that are otherwise lost. Specifically, intra-cortical edges are weighed by they 3D Euclidean distance; white matter edges are weighed by the distance along the respective DTI fiber path, divided by a factor of 200. This value is chosen to reflect the myelination of white matter fibers, which is known to allow cortico-cortical activity to propagate at speeds ∼ 200 times greater in comparison with intra-cortical propagation [21]. Resting-state BOLD fMRI timecourses of the subject were minimally preprocessed (coregistration, motion correction), resampled on the subject connectome graph, and demeaned.

### Spatiotemporal observables in numerical simulations

In numerical simulations, the spatial power spectrum is obtained following its standard definition, as the temporal mean of the squared graph Fourier transform of activity fluctuations about a steady state. The temporal power spectrum is estimated with the periodogram method, as implemented in the Scipy Python package. The correlation matrix is obtained by normalizing the covariance matrix of excitatory activity, which was itself estimated with the Numpy Python package.

### Code repository

All code used for analysis and simulations is available for use and review at https://github.com/marcoaqil/Graph-Stochastic-Wilson-Cowan-Model

## Discussion

In this work, we have presented a general approach to whole-brain neural activity modelling on unparcellated connectomes (*graph neural fields*), by combining tools from graph signal processing and neural field equations. We developed a technique to compute spatiotemporal observables (CHAOSS), and showed that a Wilson-Cowan stochastic graph neural field model can reproduce the empirically observed harmonic spectrum of resting-state BOLD signal fluctuations. Graph neural fields can address some limitations of existing modelling frameworks, and therefore represent a complementary approach resulting particularly suitable for mesoscopic-scale modelling and connectome-graph-based analyses. To discuss advantages and limitations of our approach, it is useful to contextualize it within the existing landscape of whole-brain models.

Existing whole-brain models can be broadly divided into two classes, according to whether they incorporate short-range intracortical connectivity or not. *Region-based models* only take into account long-range connectivity between dozens or few hundreds of macroscopic ROIs, whereas *surface-based models* directly incorporate short-range intra-cortical connectivity as well [26, 30]. It is furthermore possible to distinguish between *discrete* and *continuous* surface-based models. Discrete surface-based models are defined on a (highly-sampled) cortex and are therefore finite-dimensional. In several studies, region-based and discrete surface-based models are collectively referred to as *networks of neural masses* [17, 31, *32]*. *Continuous surface-based models are better known as neural field models* and are defined on the entire cortex and are thus infinite-dimensional [23, 26, 33]. Mathematically, discrete surface-based models are finite-dimensional systems of ordinary differential equations, whereas neural field models are partial integro-differential equations.

Region-based models are constructed by parcellating the cortex into a number of regions-of-interest (ROIs), placing a local model in each ROI, and connecting them according to a given connectome (see [2, 17, 32] for reviews). The ROIs are usually obtained from structural or functional cortical atlases and the number of ROIs is in the order of a hundred or less. Connectome-based mass models are characterized by the type of local models and how they are connected i.e. if the connections are weighted or not, excitatory or inhibitory, and if transmission delays are incorporated. A wide variety of local models has been used in the literature, including neural mass models, self-sustained oscillators, chaotic deterministic systems, circuits of spiking neurons, normal-form bifurcation models, rate models, and density models [2, 17, 33]. Region-based models have proven valuable in understanding varies aspects of large-scale cortical dynamics and their roles in cognitive and perceptual processing, but they are limited in one important respect: they do not allow studying the spatiotemporal organization of cortical activity on scales smaller than several squared centimeters and their effects on large-scale pattern formation. This is due to the fact that the dynamics within ROIs are modeled by a single model without spatial extent. This prevents studying the mechanisms underlying a large class of cortical activity patterns that have been observed in experiments, including traveling and spiral waves, sink-source-dynamics as well as their role in shaping macroscopic dynamics. This is a significant limitation, particularly because the role of mesoscopic spatiotemporal dynamics in cognitive and perceptual processing is increasingly being recognized and experimentally studied [34,35]. Graph neural fields present the advantage of allowing explicit modelling of activity propagation dynamics with spatiotemporal convolutions and graph differential equations on mesoscopic-resolution connectomes, thereby overcoming this limitation.

Whole-brain models that incorporate short-range connectivity are referred to as *surface-based* because they can are defined either on high-resolution surface-based representations of the cortex [30, 36, 37] or on the entire cortex viewed as a continuous medium. We will refer to these types of models as *discrete* and *continuous* surface-based models, the latter of which are known as *neural field models* [23, 26, 38, 39]. Numerically simulating discrete surface-based models is much more computationally demanding than simulating region-based models as the former typically have dimensions that are one to two orders higher than those of the latter. Numerically simulating neural field models is even more demanding and requires heavy numerical integration in combination with advanced analytical techniques [40]. Moreover, simulating neural field models requires special preparation of cortical meshes to ensure accuracy and numerical stability. [36, 37, 41–43]. In this context, graph neural fields have the advantage of being implementable natively on multimodal structural connectomes obtained from MRI and DTI, thereby minimizing anatomical approximations. Graph neural fields can naturally take into account important physical properties such as cortical folding, hemispheric asymmetries, non-homogeneous structural connectivity, and white matter projections, with a minimal amount of computational power. Furthermore, the cortex in graph neural field need not be a flat or spherical bidimensional manifold. The approach can be straightforwardly extended to include cortical thickness, allowing activity to propagate not only tangentially, but also perpendicularly to the cortical surface. Cortical layers can already be distinguished with high-field and ultra-high field functional fMRI, and are thought to subserve different functions [44]. The ability of graph neural fields to account for cortical thickness in dynamical models of neural activity is therefore a promising property for future development [45].

Our approach presents several limitations. First, some of the analytic results presented here (CHAOSS) rely on the model parameters being space-independent, that is the model parameters are assumed to be the same throughout the cortex. This assumption has the advantage of allowing mathematical analyses that, unlike numerical simulations, are virtually “infinitely” scalable with little computational cost, and was also used in previous studies of continuous neural fields [18]. However, there are more biophysically realistic models that require space-dependent parameters. For example, some recent neural mass network models incorporate neuronal receptors and their densities, which are known to vary across the cortex [46–48]. There are several ways in which it might be possible to overcome this limitation. We remark that it is only our analytic approach that requires space-independent parameters; numerical simulations of graph neural fields could be carried out also with space-dependent parameters (of course, such simulations would be more computationally demanding than their counterparts with space-independent parameters). To preserve analytic tractability while characterizing regional differences, one could attempt to absorb all the relevant space-dependent information into the graph Laplacian. Similarly to the idea of differentially weighing white matter edges to account for myelination, one might weigh differentially graph edges within specific ROIs or specific subsets of vertices. A hybrid approach of space-dependent numerical simulations and space-independent analysis (by averaging values of space-dependent parameters) could be another way to address this issue. Second, we have restricted our approach to spatially symmetric kernels. In some special cases, asymmetric kernels may be practically obtained by introducing suitable asymmetries in the graph edges. For example, consider a grid graph in two dimensions, with additional edges connecting bottom-left and top-right vertices of each square in the grid. Because of the broken lattice symmetry, a Gaussian kernel on this non-grid graph will behave like a spatially elliptic Gaussian, angled at 45 degrees. This is analogous to modelling a spatially asymmetric diffusion process on the graph. Third, another limitation is the use of an undirected and time-independent connectome graph. For maximal generality and biophysical realism, one might want to study a directed, or even time-dependent (plastic) structural connectome: it is not clear at present if and how this could be implemented in the framework of graph neural fields.

Immediate applications of graph neural fields can be found in the comparison of harmonic spectra, functional connectivity and coherence matrix with single-subject empirical data obtained from different neuroimaging modalities such as fMRI and MEG, as well as different conditions, for example health, pathology, and neuropharmacologically altered states of consciousness [22]. For example, investigating the effects of a reduced myelination speed factor, or pruned white-matter fibers, could be an interesting approach to modelling the effects of pathological or age-related structural alterations on the dynamics of functional activity. Other applications include the implementation of more biophysically realistic models, potentially including space-dependent parameters, and the use of a cortical connectome that includes cortical thickness, accounting for activity propagation across layers perpendicularly to the surface. Aside from whole-brain modelling, graph neural fields may also be used for modelling specific ROIs and stimulus-evoked brain activity. In particular, because of the known retinotopic mapping between visual stimuli and neural activity, the visual cortex presents itself as a very interesting ROI for such developments [49]. Moving beyond neural populations and even the human brain, we note that the graph Laplacian may also be used to implement single-neuron models directly on the full connectome graphs of simple organisms, such as *C*. *Elegans*, whose connectome has been experimentally mapped at the single-neuron level.

## Conclusion

In this study we described a class of whole-brain neural activity models which we refer to as *graph neural fields*, and showed that they can be used to investigate properties of brain activity measured with neuroimaging methods. The formulation of graph neural fields relies on existing concepts from the field of graph signal processing, namely the graph Laplacian operator and graph filtering, and modelling concepts such as neural field equations. This framework allows inclusion of realistic anatomical features, analytic predictions of harmonic-temporal power spectra, correlation, and coherence matrices (CHAOSS), and efficient numerical simulations. We illustrated the practical use of the framework by reproducing the harmonic spectrum of resting-state BOLD fMRI with a stochastic Wilson-Cowan graph neural field model. Future work could build on the methods and results presented here, both from theoretical and applied standpoints.

## Supporting information

**S1 Table.** Parameter set for 1D analysis and simulations. This parameter set was obtained by a qualitative comparison of the Wilson-Cowan model’s harmonic and temporal spectra with empirical data, and used to illustrate how graph properties affect neural field dynamics in one dimension.

| Parameter | Value | Units (S.I) |
| --- | --- | --- |
| $\tau_E$ | $2.415 \cdot 10^{-1}$ | s |
| $\tau_I$ | $5.227 \cdot 10^{-1}$ | s |
| $\sigma_{EE}$ | $4.577 \cdot 10^{-3}$ | m |
| $\sigma_{IE}$ | $1.704 \cdot 10^{-3}$ | m |
| $\sigma_{EI}$ | $9.377 \cdot 10^{-3}$ | m |
| $\sigma_{II}$ | $2.239 \cdot 10^{-1}$ | m |
| $d_E$ | $10^2$ | - |
| $d_I$ | 9.430 | - |
| $\alpha_{EE}$ | $6.886 \cdot 10^2$ | - |
| $\alpha_{IE}$ | $7.903 \cdot 10^2$ | - |
| $\alpha_{EI}$ | $9.972 \cdot 10^2$ | - |
| $\alpha_{II}$ | $1.223 \cdot 10^2$ | - |
| $P$ | 3.824 | - |
| $Q$ | 7.120 | - |
| $\sigma$ | $10^{-7}$ | - |

**S2 Table.** Parameter set for connectome-wide analysis and simulations. This parameter set was obtained by quantiatively fitting the Wilson-Cowan model’s harmonic power spectrum to that of resting-state fMRI data, and used for all connectome-wide analysis and numerical simulations.

| Parameter | Value | Units (S.I) |
| --- | --- | --- |
| $\tau_E$ | $2.024 \cdot 10^{-1}$ | s |
| $\tau_I$ | $2.346 \cdot 10^{-1}$ | s |
| $\sigma_{EE}$ | $1.611 \cdot 10^{-2}$ | m |
| $\sigma_{IE}$ | $2.022 \cdot 10^{-3}$ | m |
| $\sigma_{EI}$ | $6.698 \cdot 10^{-2}$ | m |
| $\sigma_{II}$ | $9.149 \cdot 10^{-2}$ | m |
| $d_E$ | $2.718 \cdot 10^1$ | - |
| $d_I$ | 1.240 | - |
| $\alpha_{EE}$ | $1.487 \cdot 10^2$ | - |
| $\alpha_{IE}$ | $2.191 \cdot 10^2$ | - |
| $\alpha_{EI}$ | $2.620 \cdot 10^2$ | - |
| $\alpha_{II}$ | $1.614 \cdot 10^2$ | - |
| $P$ | $2.235 \cdot 10^1$ | - |
| $Q$ | 8.450 | - |
| $\sigma$ | $10^{-7}$ | - |

**S3 Fig.**
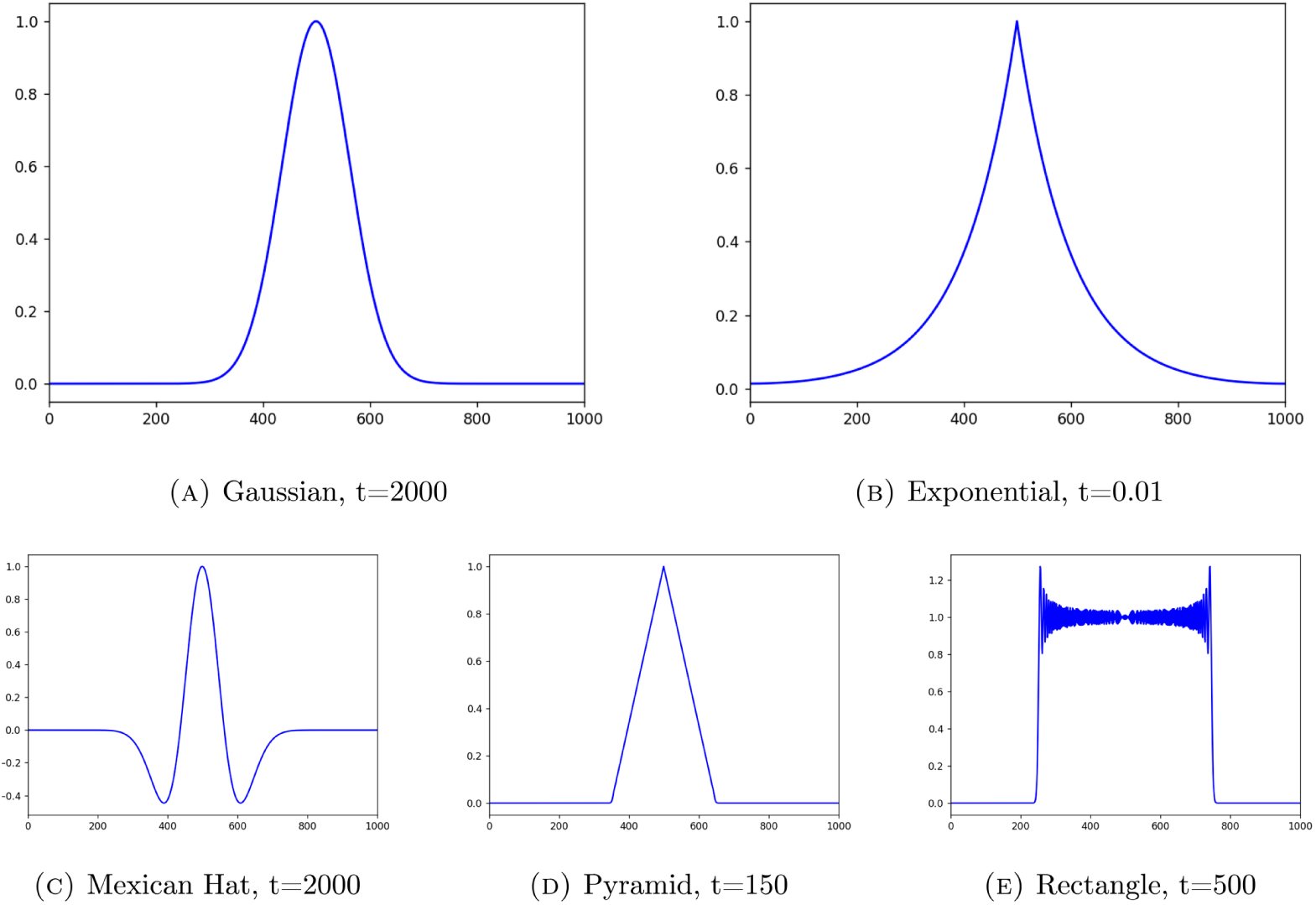
Spatial convolutions on 1-dimensional graphs. To illustrate spatial convolution on graphs, we apply different spatial convolution filters from Table 2 to an impulse function centered on the mid-node of a one-dimensional grid-graph with spacing *h* = 1 units. The resulting functions, normalized to have unit amplitude, show the shapes of the graph kernels. Note that the rectangular kernel convolution operator in Panel (E) exhibits the *Gibbs phenomenon* [50], which is a known feature of finite Fourier representations of functions with jump discontinuities. Solutions to this problem have been offered [51], but they are beyond the scope of the current work. Thus, we suggest avoiding spatial kernels with jump discontinuities in the context of graph neural fields. As a side remark, we also note that for large datasets (for example natural images databases), it might be computationally advantageous to apply convolutions with symmetric kernels through graph filters, rather than with standard discrete convolution methods. Spatial convolutions on graphs become linear matrix-vector products, which are highly optimized and easily parallelizable operations; the bulk of the computational cost for graph convolutions consists in the initial computation of the filter itself, which has to be performed only once per kernel.

## Acknowledgments

We would like to thank Thomas Yeo and Ruby Kong for providing the mapping between HCP 32k and 10k vertices and Daniele Avitabile for valuable discussions. MLK is supported by the ERC Consolidator Grant: CAREGIVING (n. 615539), Center for Music in the Brain, funded by the Danish National Research Foundation (DNRF117), and Centre for Eudaimonia and Human Flourishing funded by the Pettit and Carlsberg Foundations. RH is supported by the NWO-Wiskundeclusters grant nr. 613.009.105.

## Footnotes

1 that is, graphs equipped with a suitable distance metric

2 These assumptions are required because generally space-dependent parameters would be expressed by a non-constant diagonal matrix, that would not commute with the graph Fourier transform *U*^*T*^.

3 Eigenmode-dependent parameters would be expressed by a diagonal matrix in the graph Fourier domain, therefore by definition commuting with *U*^*T*^.

4 One could also write ∇ = [∂_1_, …, ∂_*N*_] for the “graph-gradient”, and then define Δ = − ∇.*·*∇. The minus sign arises to maintain the analogy with the continuous Laplacian.

5 Alternatively, for the case of space-dependent parameters, observables may be computed from numerical simulations

